# CcBHLA: pan-specific peptide–HLA class I binding prediction via Convolutional and BiLSTM features

**DOI:** 10.1101/2023.04.24.538196

**Authors:** Yejian Wu, Lujing Cao, Zhipeng Wu, Xinyi Wu, Xinqiao Wang, Hongliang Duan

## Abstract

Human major histocompatibility complex (MHC) proteins are encoded by the human leukocyte antigen (HLA) gene complex. When exogenous peptide fragments form peptide-HLA (pHLA) complexes with HLA molecules on the outer surface of cells, they can be recognized by T cells and trigger an immune response. Therefore, determining whether an HLA molecule can bind to a given peptide can improve the efficiency of vaccine design and facilitate the development of immunotherapy. This paper regards peptide fragments as natural language, we combine textCNN and BiLSTM to build a deep neural network model to encode the sequence features of HLA and peptides. Results on independent and external test datasets demonstrate that our CcBHLA model outperforms the state-of-the-art known methods in detecting HLA class I binding peptides. And the method is not limited by the HLA class I allele and the length of the peptide fragment. Users can download the model for binding peptide screening or retrain the model with private data on github (https://github.com/hongliangduan/CcBHLA-pan-specific-peptide-HLA-class-I-binding-prediction-via-Convolutional-and-BiLSTM-features.git).

## Introduction

The major histocompatibility complex (MHC) is a group of genes encoding cell surface proteins, which is of great significance for the adaptive immune system to recognize exogenous molecules and respond accordingly. Human MHC proteins are encoded by the human leukocyte antigen (HLA) gene complex. In general, exogenous molecules are first processed into peptide fragments and then presented to HLA molecules on the extracellular surface to form a peptide-HLA (pHLA) complex, which is recognized by T cells to trigger a powerful immune response[1]. Thus, the binding of peptides to HLA is an essential prerequisite for efficacious T-cell recognition[2].

HLA molecules can be broadly classified into HLA I class and HLA II class based on their function, molecular structure and distribution. HLA class I is an endogenous antigen-presenting protein, mainly binding short peptides with a length of 8-12 amino acids, which are usually derived from proteasome-mediated intracellular protein degradation. CD8+ T cells primarily recognize these pHLA complexes presented on the cell surface. HLA class II are exogenous antigen-presenting proteins that tend to bind peptides 12-20 amino acids in length. These pHLA complexes occur on professional antigen-presenting cells (APCs), such as macrophages, dendritic cells, and B cells, and are recognized by CD4+ T cells[3, 4]. Due to the ubiquity of HLA class I, it has been more widely concerned in previous biomedical research[5, 6]. Understanding whether HLA molecules can bind to a given peptide will help workers better explore the mechanism of immune recognition, speed up the identification of new antigens and improve the efficiency of vaccine design[7]. Traditional research methods usually use gold standard methods such as enzyme-linked immunosorbent spot, competitive binding assays and the direct binding assay. However, these wet lab validation methods are financially burdensome and inefficient.

With the development of artificial intelligence, a large number of tools for inferring potential binding affinities between HLAs and peptides have been developed in the past few years. These methods can be roughly divided into the following three categories. The first is the structure-based approach[8] which relies on the analysis of the binding structure of HLAs and peptides, such as a statistical energy function based on residual[9], quantitative structure-activity relationship analysis[10] and docking[11–14]. Although this method helps researchers to understand the binding of HLA and peptide more intuitively at the structural level, the difficulty in obtaining shared and available crystal structures restricts the improvement of the method in terms of accuracy, scope and prediction speed. The second is a function-based scoring method, which calculates the features of peptide sequences to generate a scoring matrix for specific HLA allotype for subsequent pHLA binding prediction. The most representative method is Anthem[15] proposed by Mei et al. Unlike previous methods that only used one or two scoring functions[16–19], Anthem applies five scoring functions, including amino acid frequency, webLogo-based sequence conservation, two types of PSSMs[18], and BLOSUM62[20], comprehensively encode each peptide sequence. The model is then trained by integrating the results of five scoring functions in the set and achieves better performance. Finally, there is the deep learning approach, which has shown powerful capabilities in extracting high-dimensional features of peptide sequences through multilayer neural networks. Especially in recent years, the large-scale availability of immune peptide population data generated by applying mass spectrometry[21, 22] provides a growing opportunity for the development of data-driven deep learning models for pHLA binding prediction. As far as we know, the most advanced deep learning method for predicting the binding affinity between HLAs and peptides is TransPHLA constructed by Chu et al.[23] The core of TransPHLA is based on self-attention, and extracts sequences features of peptides and HLA through the embedding block, the encoder block, the feature optimization block, and the projection block. Finally, the features are passed through the fully connected layer to calculate the binding affinity. The TransPHLA has higher performance and efficiency, both compared to specific and pan-specific methods.

In previous studies on peptides using deep learning, the convolutional neural network (CNN) has been widely used because of its ability to extract key features of peptide sequences, and can achieve excellent performance[24–27]. However, CNN ignores the position information when extracting sequence features. It is well known that the position of amino acids in a peptide is essential information, which affects and even determines the spatial structure and function of a peptide. Models designed to process natural language generally have the ability to extract contextual information and have been applied to peptide processing[28–30]. Therefore, we explore whether CNN can be combined with a natural language processing model to bring better performance.

In this paper, we construct a novel deep learning-based model called CcBHLA (Convolutional Neural Networks Combined with Bilstm for HLA Class I Binding Prediction) to accurately predict the binding affinity between HLA Class I and peptides. We set the kernel_size of CNN to 2, 3, and 4 respectively, which is similar to extracting n-gram features during the movement process, while BiLSTM pays more attention to the context information of the peptide sequence. Our method is pan-specific and is not limited by the length of the predicted peptide. The peptide and HLA sequences are encoded as onehot vectors without any additional information (such as scoring functions). Compared with the state-of-the-art TransPHLA model, our method achieves superior performance on all metrics. In addition, we further explored the impact of the sequence information of HLA on the prediction performance. We upload the model’s code to github, where users can download it for free to make testable predictions, or retrain the model on their data.

## Materials and Methods

### Data

In this study, all data were derived from TransPHLA, which can be downloaded from https://github.com/a96123155/TransPHLA-AOMP/tree/master/Dataset. The pHLA binding data (positive data) in the data were collected by Anthem, while negative data were randomly selected from source proteins in IEDB by TransPHLA referring to previous studies[31–33]. The positive and negative data of the same peptide length under each HLA allele are equal to warrant the balance of positive and negative samples in the constructed data set.

The data set is divided into three categories: training set, independent test set and external test set. The training set is used for model training and selection, the independent test set is used for model performance evaluation, and the external test set is used for model performance comparison. The data of both the training set and the independent test set were from (1) four public databases of IEDB[34], EPIMHC[35], MHCBN[36] and SYFPEITHI[37], (2) data identified by mass spectrometry in previously published studies (3) training data sets collected by other pHLA binding prediction tools. Therefore, the distributions of the training set and the independent test set are very cognate, but there is no overlap with each other. To avoid inflated model performance due to similar data distributions, the experimentally validated data in Anthem is used as an external test set for a fair comparison. Notably, since the typical length range for HLA-I peptide binding is 8–14 amino acids[38], only peptides within this length range were selected for each HLA-I allotype.

Following TransPHLA’s data splitting, we performed five-fold cross-validation (CV), in which the training set was randomly divided into five equal parts, four of which were used as training data, and the remaining one was used as a validation set to evaluate the training of the model. Results for all metrics in our paper are the average of five model results.

### Data encoding

Since the spatial structure and function of the peptide are determined by its primary structure, the type and arrangement of amino acids in a peptide contain rich feature information, which can be mined by a neural network to facilitate the prediction of pHLA binding affinity. All peptides in the data consist of 20 amino acids. According to the official IUPAC-based amino acid single-letter notation[39, 40], the amino acid alphabet is defined as ={A, R, N, D, B, C, E, Q, Z, G, H, I, L, K, M, F, P, O, S, U, T, W, Y, V, X}. The input data for our prediction framework CcBHLA includes peptide and HLA sequences. First, the peptide and HLA sequences are joined with a lowercase “z” that does not exist in the amino acid alphabet to form the complete pHLA sequence. Then use single-character embedding to create a unique embedding for each amino acid in pHLA and the connection symbol ‘z’. The dimension of the embedding is defined as 21×L, where L is the length of the pHLA sequence. Taking the pHLA sequence ‘QLKEYLFYzYFAMYQENMAHTDANTLYIIYRDYTWVARVYRGY’ as an example, its length is 43. Finally, the pHLA sequence embeddings are padded to a maximum length of 50 to accommodate variable input lengths. In addition, we do not add any additional information such as scoring matrix to avoid possible wrong features.

## Model

### BiLSTM

Long Short-Term Memory (LSTM) is a type of Recurrent Neural Network (RNN)[41]. Owing to its design characteristics, LSTM is very suitable for modeling time series data, such as text data. LSTM can learn which information to remember and which information to forget during the training process, so it can better capture longer-distance dependencies in text. When LSTM is used to model sequences, it is impossible to encode information from back to front, thus deriving Bi-directional Long Short-Term Memory (BiLSTM)[42]. BiLSTM is a combination of forward LSTM and backward LSTM, which can well capture bidirectional semantic dependencies and is often used to model contextual information in natural language processing tasks.

In our task, the amino acid single-character embedding matrix of pHLA with a size of 50×21 is first fed into the Embedding layer to amplify some features or separate some redundant features. The vector matrix output by the Embedding layer is input to BiLSTM, and the output of BiLSTM is concatenated from the vector matrix obtained by processing the forward and backward sequences.

### CNN

Convolutional Neural Network (CNN) is a type of Feedforward Neural Network that includes convolution calculations and has a deep structure[43]. It is one of the representative algorithms for deep learning. The convolutional neural network has the ability of representation learning, and can perform shift-invariant classification on the input information according to its hierarchical structure, so it is also called “Shift-Invariant Artificial Neural Networks (SIANN)”.

CNNs were originally proposed for image recognition tasks[44] and have flourished in computer vision. In 2014, Yoon Kim made some changes to the input layer of CNN and proposed textCNN for text classification[45]. This is the first time CNNs have been applied to natural language and have been found to be very effective in processing general sequence data. Compared with the CNN in the image field, the biggest difference of textCNN is the input data: the image is two-dimensional data, and the image convolution kernel slides from left to right and from top to bottom to carry out feature extraction. Natural language is one-dimensional data. Although two-dimensional vector is generated by word embedding, it is meaningless to convolve the word vector from left to right, so the convolution kernel of textCNN only slides from top to extract features.

The CNN architecture we apply in this paper consists of convolutional layers and max pooling layers. The convolution kernel of the convolutional layer is set to three sizes of 2, 3, and 4. In the process of moving from top to bottom, the features of different sizes amino acid phrases are extracted, as shown in Figure 1. Each convolutional layer uses a Rectified Linear Unit (’ReLU’) as the activation function in the activation layer. The max pooling layer is connected after the convolutional layer to reduce information redundancy while retaining the features, thereby reducing overfitting and speeding up calculations. And finally combine the outputs of the three max pooling layers to form the final feature.

**Figure 1:**
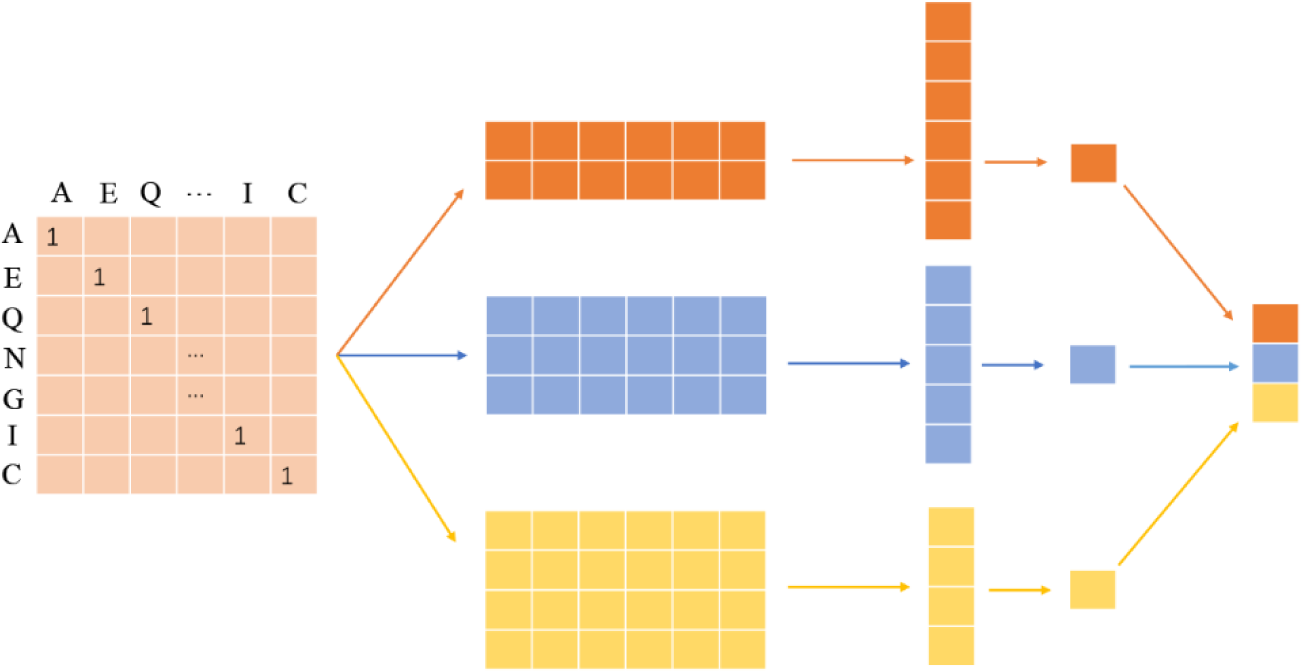
Schematic diagram of textCNN feature extraction.

### CcBHLA

The overall architecture of the model is shown in Figure 2. The input of the CcBHLA network is the amino acid character sequence of pHLA. After single-character embedding, a 50×21 one-hot matrix is obtained. The one-hot matrix is fed into the embedding layer, and each amino acid in pHLA is represented by 128 dimensional vector space. The output of embedding layer is a 50×128 vector matrix. The vector matrix is then fed into BiLSTM and returns a feature vector matrix of the same size as the input. Afterwards, the BiLSTM feature vector matrix will be input into the three CNNs with convolution kernels of 2, 3, and 4 respectively, and concatenate the output vectors of the max pooling layers of the three CNNs to obtain the CNN feature vector matrix. Finally, the feature vector matrix of CNN and BiLSTM is concatenated to obtain the final feature vector matrix of pHLA. The feature vector matrix is then connected with two dense layers whose activation function is ‘ ReLU ‘, and the dense layer is fully connected to the logistic regression output unit for prediction. We set 50% Dropout before each dense layer, so that a certain proportion of units are randomly discarded during the training process, which plays a role of regularization and prevents overfitting[46].

**Figure 2:**
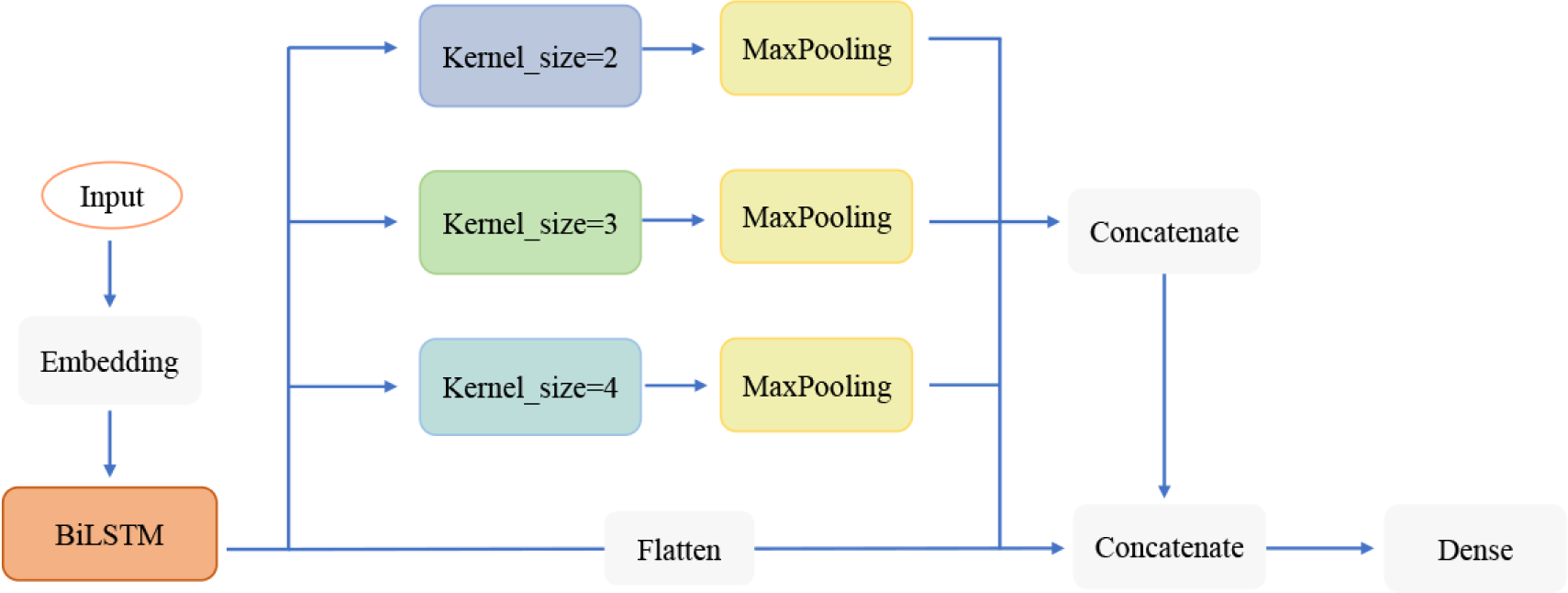
Schematic diagram of the CcBHLA model architecture. Performance evaluation metrics.

In this paper, we refer to TransPHLA and evaluate the performance of CcBHLA using metrics including accuracy (ACC), Matthews correlation coefficient (MCC), F1, and Area Under the Receiver Operating Characteristic Curve (ROC_AUC). The calculation of the above metrics is described in equations 1-4, where TP stands for true positive, TN stands for true negative, FP stands for false positive, and FN stands for false negative.

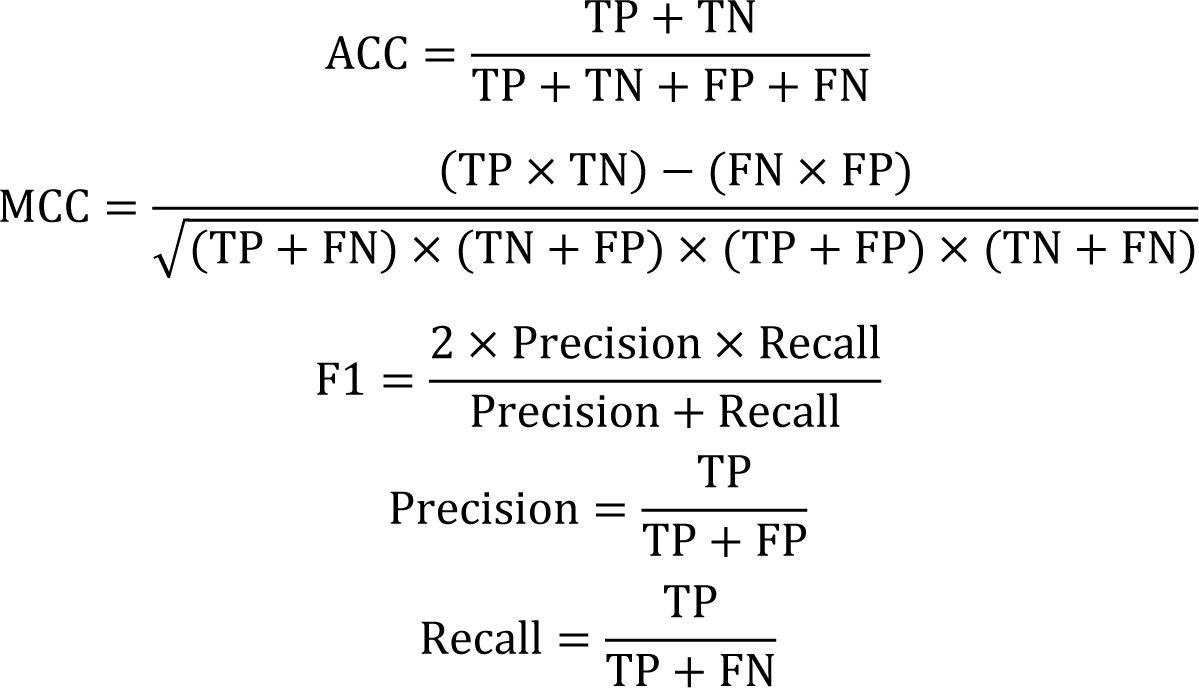

Except for MCC, which takes values from -1 to 1, the other metrics take values from 0 to 1. The higher the value of the metrics, the better the performance of the model.

## Result

### Performance comparison on the test set

Because the TransPHLA model was compared with 14 previous pHLA binding prediction methods, including nine baseline methods from IEDB (ANN[47], Consensus[48], NetMHCcons[49], NetMHCpan_BA[33], NetMHCstabpan[50], PickPocket[19], CombLib[51], SMM[52] and SMMPMBEC[17]), the method recommended by IEDB (NetMHCpan_EL[33]), a state-of-the-art method released in 2021 (Anthem), and three recently published attention-based methods (ACME[53], DeepNetBim[54], and DeepAttentionPan[55]). Among them, except NetMHCpan_BA, NetMHCpan_EL and TransPHLA, other methods have different limitations on the length of HLA alleles and predicted peptides. TransPHLA not only achieves the best performance with the highest efficiency, but also addresses the limitations of many methods for HLA alleles and variable-length peptides. Therefore, we only compare the performance with TransPHLA in this study. The performance of the two methods on the independent test set is shown in Figure 3A, and our model outperforms TransPHLA on the metrics of ACC, ROC_AUC, F1 and MCC. In order avoid inflated performance due to the same distribution of the independent test set and the training set, we compared the two models on the external test set, and the results are shown in Figure 3B. Both our model and TransPHLA experienced performance degradation. But our model still outperforms TransPHLA in various metrics, proving the superiority of our model.

**Figure 3:**
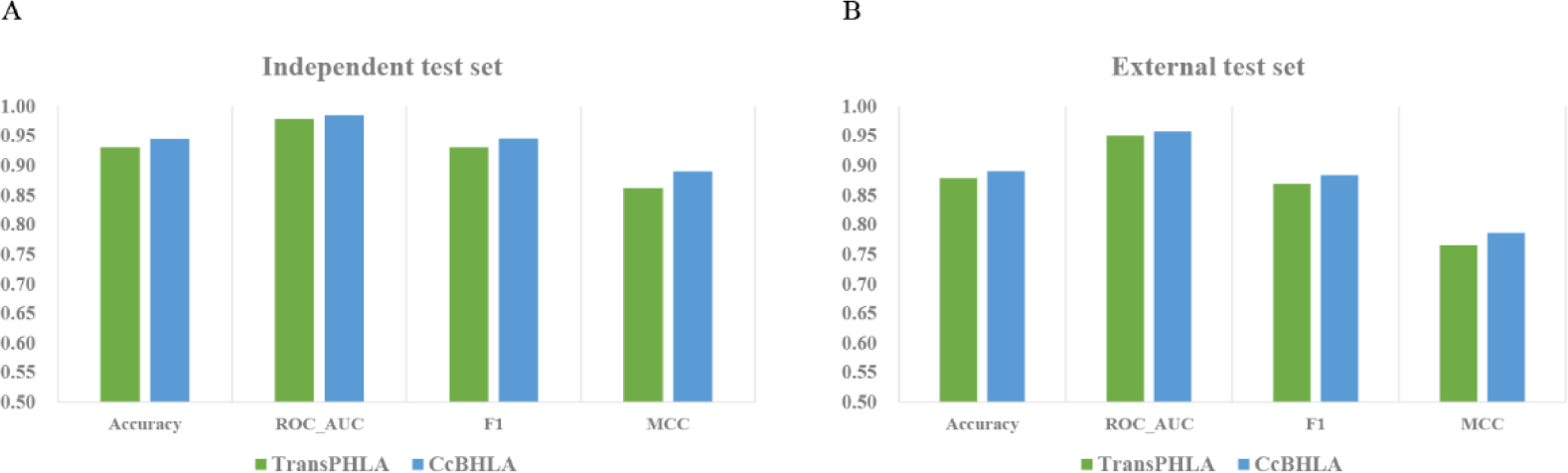
The performance of TransPHLA and CcBHLA models on the independent test set(A) and external test set(B), respectively.

### Neoantigen Screening

To demonstrate the generalization ability and generality of our model, we performed neoantigen screening using the neoantigen data collected by TransPHLA, which included 221 experimentally validated pHLA-binding peptides collected from non-small cell lung cancer, melanoma, ovarian and pancreatic cancer[56, 57]. As a major determinant of neoantigen screening, it is important to accurately predict binding of peptides to self-specific HLA molecules. The prediction results of our model and other models are shown in Figure 4, where the results of other models are derived from TransPHLA. The specific model cannot predict all neoantigen data due to length limitations. Among non-specific models, our model achieves the same prediction accuracy as TransPHLA with only 8 wrong predictions. In practical applications, it can help to speed up the screening of antigens, thereby helping the development of vaccines.

**Figure 4:**
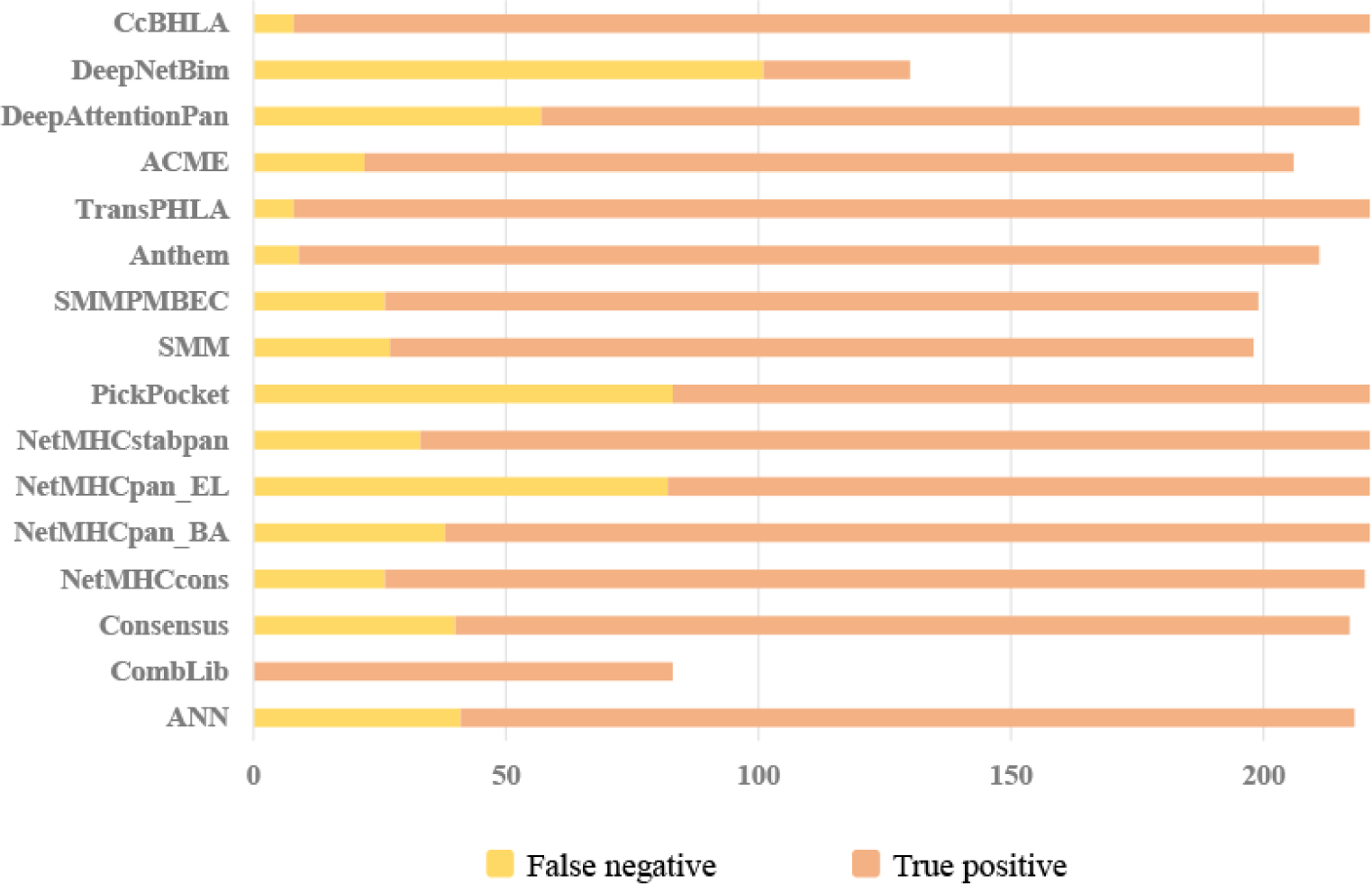
Performance of each model on 221 neoantigen screens.

### Performance comparison of each peptide length

Due to the closed conformation of the HLA class I antigen-binding groove, antigenic peptides bound by different HLA class I molecules have distinct allele-specific amino acid preferences. Recent elution studies have found that 9-mer peptides bind better to HLA molecules[58, 59]. This is likely caused by the anchor residues of the antigenic peptide corresponding to the Position2 and Position9 positions of the nine amino acid binding cores[60]. This is why peptides with a length of 9 have the most data.

The distribution of different peptide lengths in the data set is shown in Figure 5. There are great differences in the amount of data with different lengths. The number of 9-mer peptides accounts for a large proportion, while the number of 13-mer and 14-mer peptides is very small. The deep learning model relies on data-driven. The peptide length with a large amount of data can help the model learn some common feature knowledge to improve its performance on the peptide length with insufficient data. However, the limitation of the model performance due to the lack of data cannot be fundamentally solved, which will result in a wide variation in the performance of different peptide lengths. Figures 6A and 6B demonstrate the performance capabilities of our model on the independent test set and external test set for the overall and different peptide lengths, respectively.

**Figure 5:**
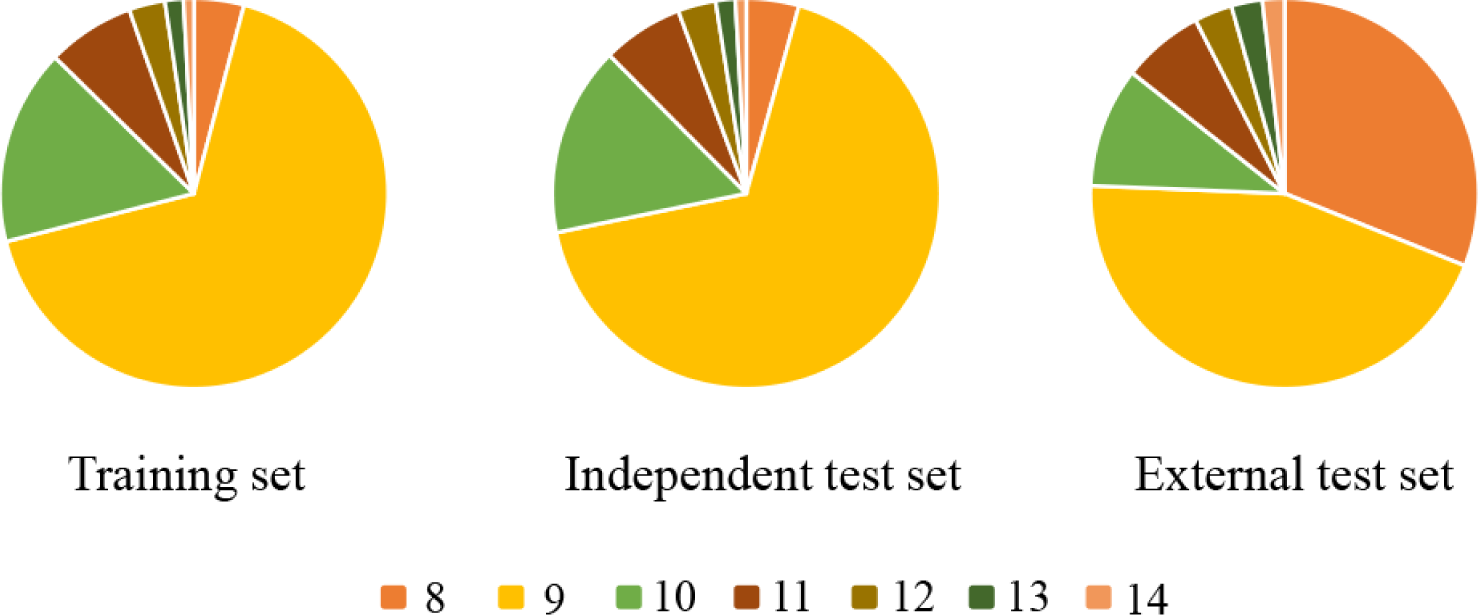
The distribution ratio of peptide lengths.

**Figure 6:**
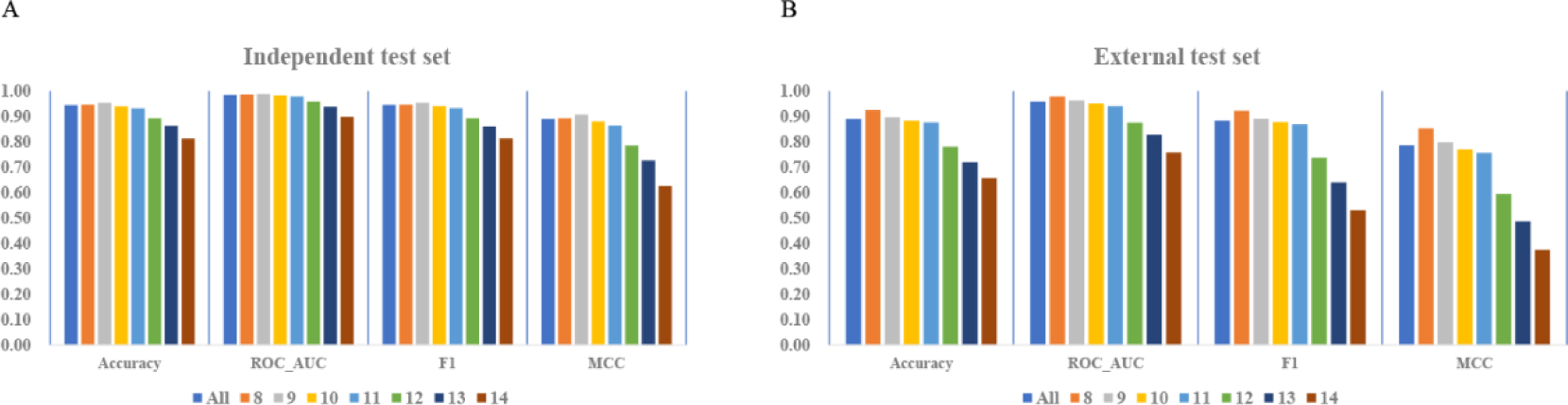
The performance of our model on the independent test set and external test set for the different peptide lengths.

Figures 7 show the performance comparisons of our model and TransPHLA on Accuracy, ROC_AUC, F1, and MCC for different peptide lengths from 8 to 14 on the independent test set, respectively. Similarly, Figures 8 is the performance comparison of the two models on the external test set. From the above figures, we can conclude that for the peptide length larger than 9, as the peptide length increases, the corresponding amount of data decreases, and the prediction performance of the two models also decreases. However, our model outperforms TransPHLA on all performance metrics per peptide length, both on an independent test set with a distribution similar to the training set, and on an external test set validated by Anthem experiments outside of the training set distribution.

**Figures 7:**
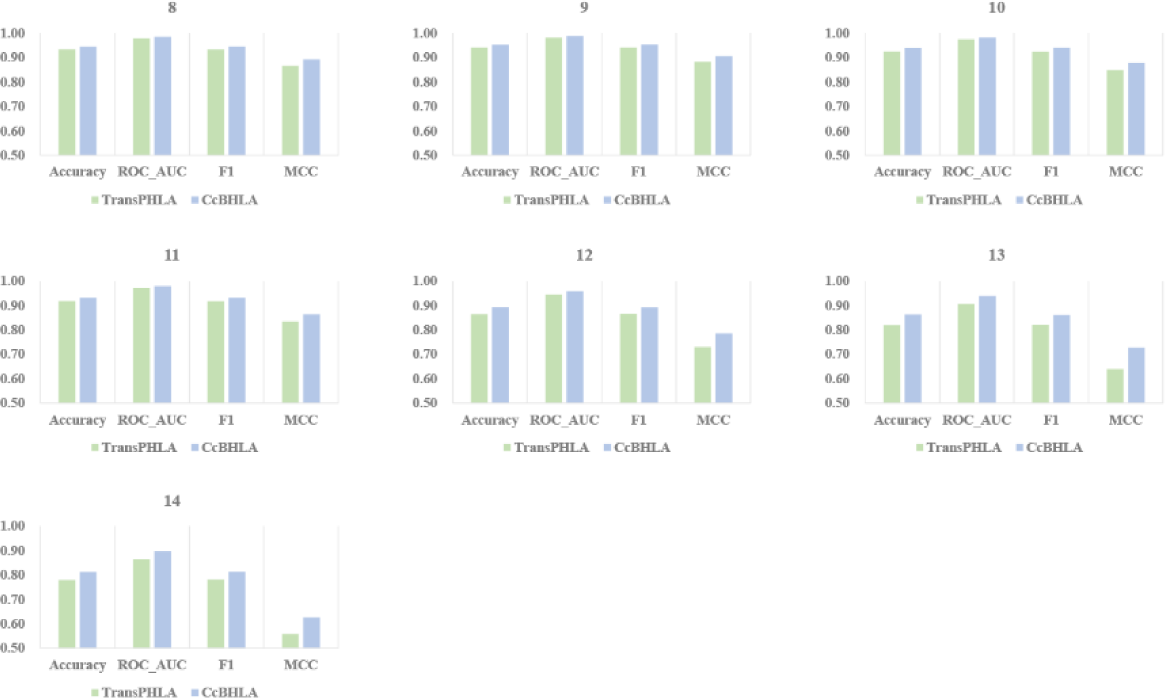
Performance comparison of CcBHLA and TransPHLA on independent test sets for different peptide lengths from 8 to 14, respectively.

**Figures 8:**
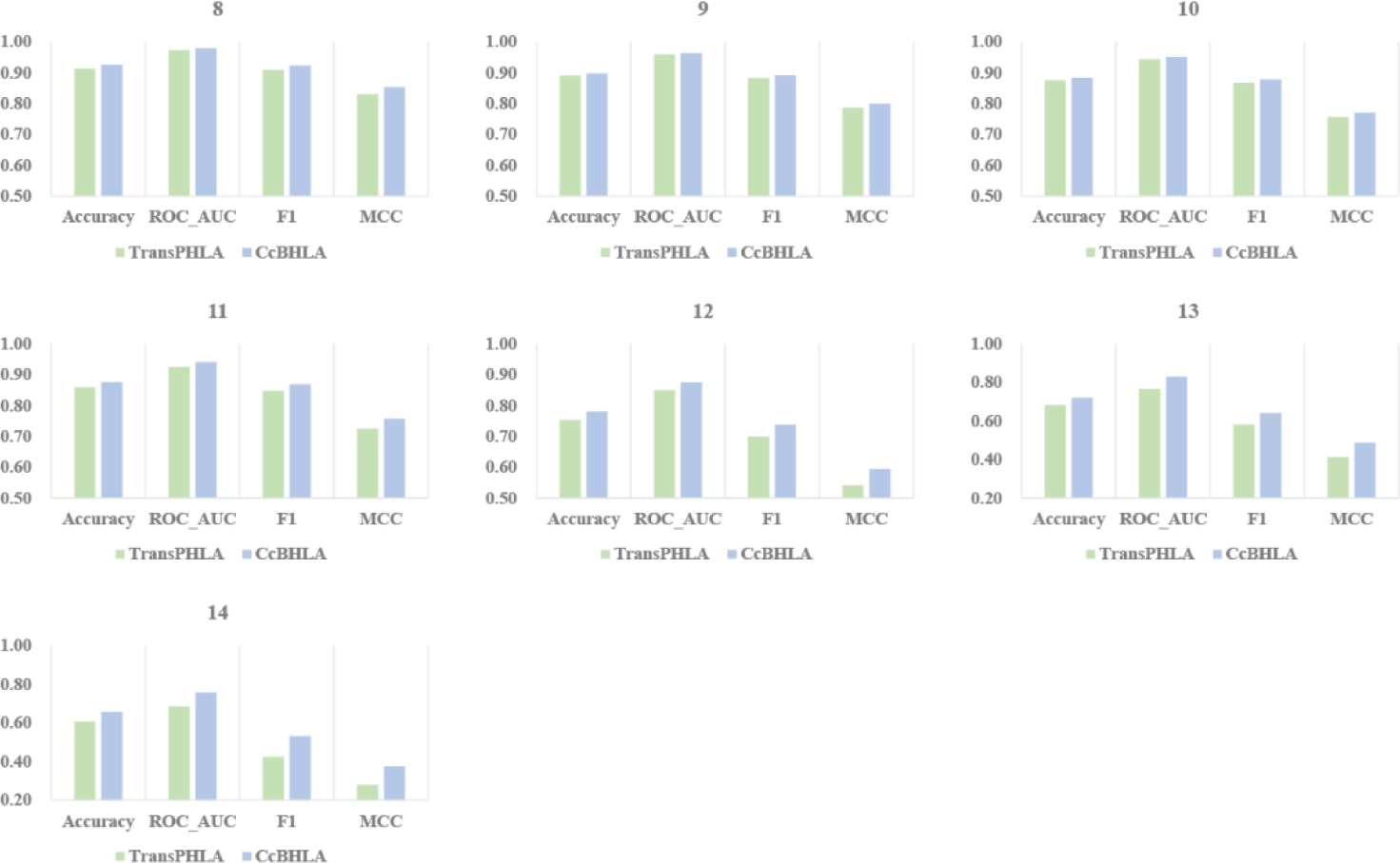
Performance comparison of CcBHLA and TransPHLA on external test sets for different peptide lengths from 8 to 14, respectively.

Besides, we evaluated the performance of the model for each peptide length under each HLA. The Accuracy, ROC_AUC, F1, and MCC of all HLAs are plotted into a violin plot, indicating the performance distribution of HLAs at a specific peptide length. In addition, from the calculation formula of MCC, we know that when any of TN and TP is 0 and any of FN and FP is 0, the denominator of MCC is 0 and cannot be calculated. Therefore, if the MCC cannot be calculated for a specific peptide length under a certain HLA, it indicates that the method is invalid for this peptide length under this HLA. The violin plots of the independent test set and the external test set are shown in Figure 9 and Figure 10, respectively. And the MCC shows that our method is effective for any peptide length under any HLAs, which is known only to be achieved by TransPHLA and Anthem.

**Figure 9:**
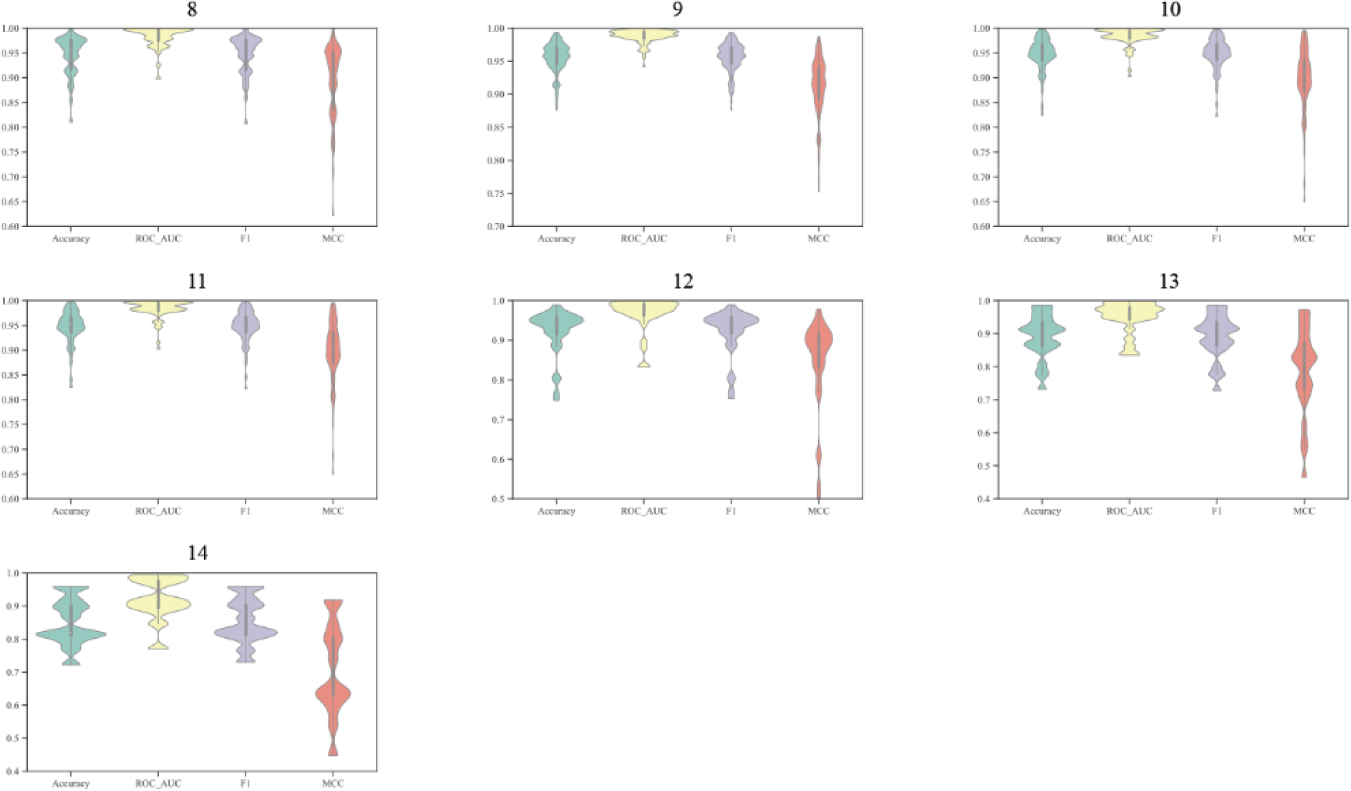
The Accuracy, ROC_AUC, F1, and MCC of the CcBHLA on the independent test set.

**Figure 10:**
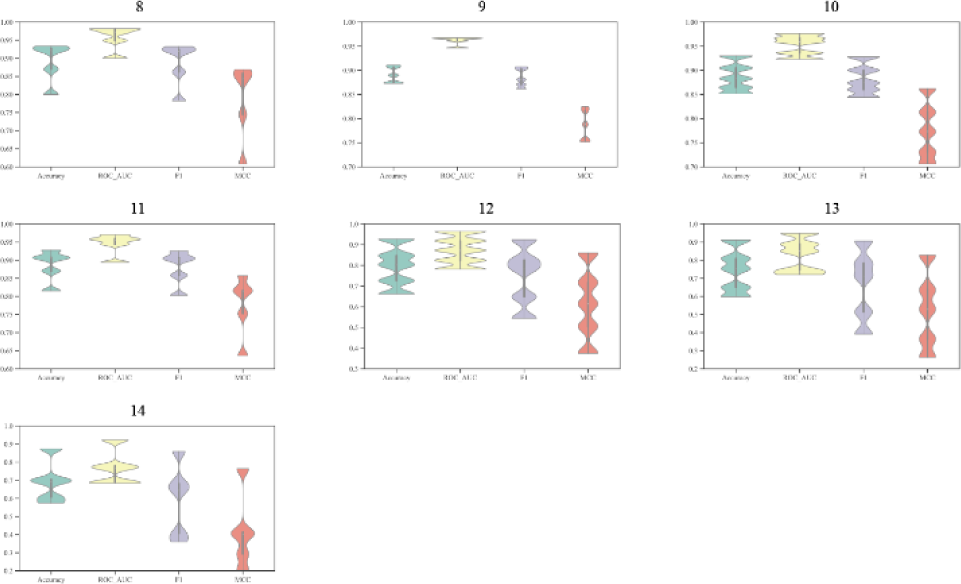
The Accuracy, ROC_AUC, F1, and MCC of the CcBHLA on the external test set.

### Hide the sequence information of HLA

The standardized nomenclature of HLA is shown in Figure 11, which is divided into three parts: gene locus, allele group, and specific HLA protein. In previous research methods, the sequence information of HLA is an essential part of the input data. It is generally believed that the interaction between amino acids in peptides and amino acids in HLA can be learned during model training. We set up an experiment in which the sequence information of HLA is removed and observe whether it will affect the predictive ability of the model. In this experiment, the sequence information of HLA was removed from the input of the model and replaced with the three-part representation of HLA shown in Figure 11, for example, the original input ‘YFAMYQENMAHTDANTLYIIYRDYTWVARVYRGY’ was replaced by ‘HLA-A,01,01’. Other model parameter settings and operation procedures remain unchanged. We re-evaluated the performance of the model on independent and external test sets, and observed how its performance varied with peptide lengths. The results are shown in Figure 12, the impact of the HLA sequence information in the input information on the performance of the model is very weak. Specific data for other assessment content can be viewed in Supporting Information.

**Figure 11:**
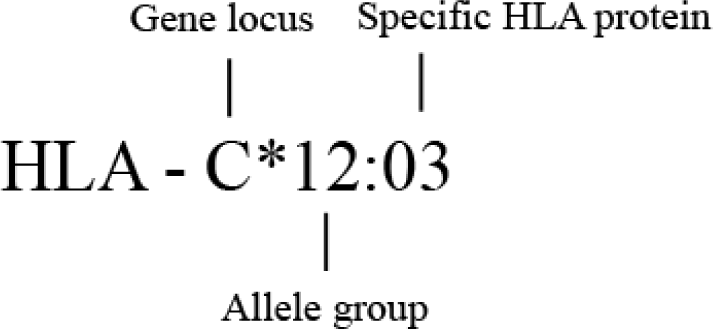
The standardized nomenclature of HLA.

**Figures 12:**
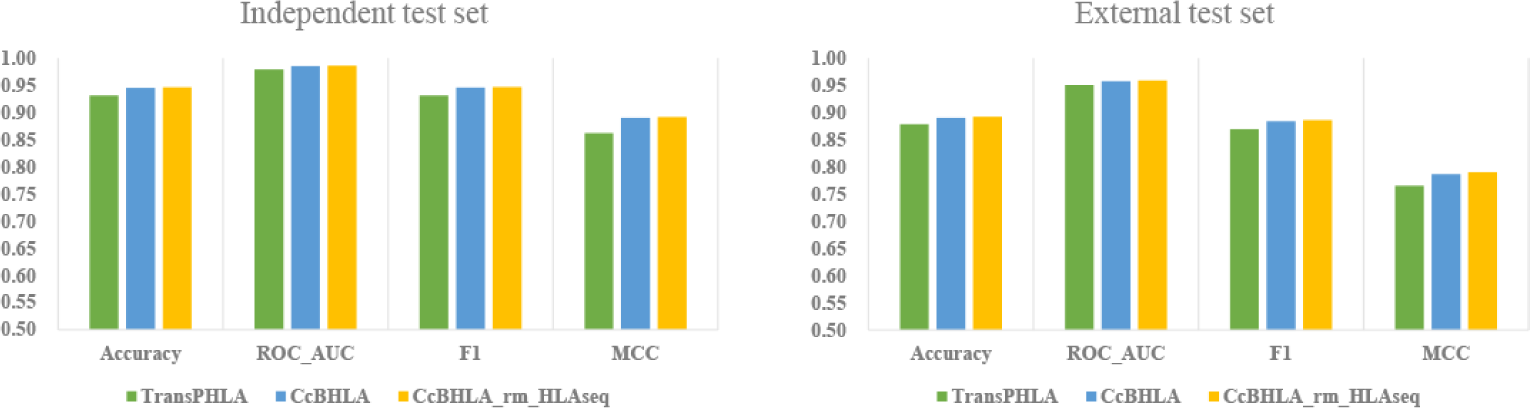
The performance of TransPHLA, CcBHLA and CcBHLA_rm_HLAseq models on the independent test set and external test set, respectively.

## Conclusion

The binding and interaction of pHLA is an important prerequisite for antigen presentation and T cell recognition. Accurately predicting the affinity of pHLA is a key step in identifying potential neoantigens and new therapeutic strategies. Many research methods have been proposed for this purpose. In this paper, we combine BiLSTM with CNN to propose a CcBHLA model for binding prediction of pHLA. The method is a pan-specific model that is not limited by HLA alleles or peptide length. We evaluate the performance of our model on an independent test set, an external test set, and screened tumor neoantigen data. By comparing with TransPHLA, the current state-of-the-art method, CcBHLA achieves excellent performance in all experimental items, and surpasses TransPHLA in the four metrics of ACC, ROC_AUC, F1 and MCC. In addition, we also found that the peptide sequence information of HLA has little impact on the performance of the model, in other words, only encoding standardized nomenclature of HLA in the data can also achieve similar performance as encoding HLA peptide sequence. By evaluating the model’s predictive performance at various peptide lengths, we determined that the performance of CcBHLA will be further improved with future data volume growth.

## Key Points

- We developed a new method for HLA-peptide binding prediction, CcBHLA, which combines the features of CNN capturing local key information and Bilstm extracting contextual information to train the model for accurate HLA-I peptide binding prediction.
- The method is pan-specific and not limited by peptide length and HLA.
- Evaluations based on independent and extended test datasets show that CcBHLA improved performance on various metrics compared to contemporary tools.

## Competing interests

No competing interest is declared.

## Data availability

The datasets are available at https://github.com/hongliangduan/CcBHLA-pan-specific-peptide-HLA-class-I-binding-prediction-via-Convolutional-and-BiLSTM-features.git and the CcBHLA package is freely available in the same GitHub repository.

## Acknowledgments

This work was supported by the National Natural Science Foundation of China (No.81903438) and Natural Science Foundation of Zhejiang Province (LD22H300004).

## Notes

### Competing Interest Statement

The authors have declared no competing interest.

